# Effects of the dynein inhibitor dynarrestin on the number of motor proteins transporting synaptic cargos

**DOI:** 10.1101/2020.02.07.934687

**Authors:** Kumiko Hayashi, Miki G. Miyamoto, Shinsuke Niwa

## Abstract

Synaptic cargo transport by kinesin and dynein in hippocampal neurons was investigated using non-invasive measurements of transport force based on non-equilibrium statistical mechanics. Although direct physical measurements such as force measurement using optical tweezers are difficult in an intracellular environment, the non-invasive estimations enabled enumerating force producing units (FPUs) carrying a cargo comprising the motor proteins generating force. The number of FPUs served as a barometer for stable and long-distance transport by multiple motors, which was then used to quantify the extent of damage to axonal transport by dynarrestin, a dynein inhibitor. We found that dynarrestin decreased the FPU for retrograde transport more than anterograde transport. In the future, these measurements may be used to quantify the damage to axonal transport resulting from neuronal diseases including Alzheimer’s, Parkinson’s, and Huntington’s diseases.

## 1. Introduction

Kinesin and dynein transport biomaterial, packaged in membrane-bound vesicles as “cargo,” along microtubules throughout eukaryotic cells(Guedes-Dias & Holzbaur, 2019; Hirokawa, Noda, Tanaka, & Niwa, 2009; Vale, 2003). These motor proteins produce their driving force by hydrolyzing adenosine triphosphate (ATP). Physical measurements of this movement, such as force-velocity curves, have been studied in single-molecule experiments(Nishiyama, Higuchi, & Yanagida, 2002; Okada, Higuchi, & Hirokawa, 2003; Schnitzer, Visscher, & Block, 2000; Tomishige, Klopfenstein, & Vale, 2002; Vale et al., 1996; Visscher, Schnitzer, & Block, 1999); however, *in vivo* mechanisms are still unclear because measuring force *in vivo* is challenging. Stokes’ law can be used to estimate *in vivo* driving force from the velocity *v* of moving cargo as the drag force through the relationship *F* = 6*πηrv*, but cargo viscosity *η* and the size *r* are difficult to measure in an intracellular environment. Optical tweezers have been used to measure force exerted on lipid droplets in cells (Jun, Tripathy, Narayanareddy, Mattson-Hoss, & Gross, 2014; Leidel, Longoria, Gutierrez, & Shubeita, 2012; Shubeita et al., 2008), but their application to submicron cargo is limited.

The *in vivo* driving force produced by motor proteins acting on intracellular cargo was investigated recently with non-invasive force measurements based on non-equilibrium statistical mechanics(Hasegawa, Sagawa, Ikeda, Okada, & Hayashi, 2019; Hayashi, 2018; Hayashi, Hasegawa, Sagawa, Tasaki, & Niwa, 2018; Hayashi, Matsumoto, Miyamoto, & Niwa, 2019; Hayashi, Tsuchizawa, Iwaki, & Okada, 2018). With the reported method, inferring the driving force from the fluctuating motion of transported cargo, the number of force producing units (FPUs), compromising motor proteins carrying a single cargo, was estimated to be 1-3 anterograde FPUs for synaptic vesicle precursor transport in motor neurons of *Caenorhabditis elegans*(Hayashi, Hasegawa, et al., 2018), 1-4 anterograde and 1-3 retrograde FPUs for endosome transport in mice dorsal ganglion neurons(Hayashi, Tsuchizawa, et al., 2018), and 1-5 retrograde FPUs for melanosome transport in zebrafish melanophores. Such multiple motor transport of cargos has been observed in force measurement experiments using optical tweezers(Leidel et al., 2012; Rai et al., 2016; Shubeita et al., 2008), in biochemical measurements(Hendricks et al., 2010), and in the analysis of transport velocity(Chowdary et al., 2015; Hill, Plaza, Bonin, & Holzwarth, 2004; Levi, Serpinskaya, Gratton, & Gelfand, 2006; Shtridelman, Cahyuti, Townsend, DeWitt, & Macosko, 2008), and is essential to understand *in vivo* transport mechanisms.

In this study, we applied the non-invasive force measurement method developed for the above systems to synaptic cargo transport in mice hippocampal neurons, noting that synaptic proteins are produced and packed in cargo vesicles in the cell body of neurons, after which the cargo vesicles are transported in an anterograde manner to the axon terminus by KIF1A and KIF1Bβ, members of the kinesin-3 family, and in a retrograde manner by cytoplasmic dynein in mammalian neurons(Hirokawa et al., 2009). Multi-motor cooperativity contributes to the stable and long-distance transport of materials along the axons to maintain neuronal activity. Additionally, the number of FPUs hauling a cargo is a reasonable indicator of healthy neuronal activity. For these reasons, we investigated the number of FPUs to understand the extent of damage caused by treatment with dynarrestin, a specific small-molecule antagonist of cytoplasmic dynein(Hoing et al., 2018). The results obtained from non-invasive force measurements showed that retrograde FPUs were decreased more than anterograde FPUs in the presence of dynarrestin. Considering the difficulties associated with direct physical measurements in live non-equilibrium neurons, we anticipate that non-invasive force measurement will aid in the determination of physical aspects of synaptic cargo transport mechanisms and potentially lead to a comprehensive, quantitative understanding of the process of synapse creation at the terminal regions of neurons. Deficits at these sites are often related to neuronal diseases, such as Alzheimer’s, Parkinson’s, and Huntington’s diseases(Encalada & Goldstein, 2014; Gabrych, Lau, Niwa, & Silverman, 2019). Thus, understanding these transport pathways could aid in identifying new therapies.

## 2. Results

### 2.1 Fluctuation of constant velocity segments (CVSs) of moving cargo

The motion of synaptic cargo, including synaptotagmins labeled with green fluorescence protein (GFP)(Niwa, Takahashi, & Hirokawa, 2013), along the axon of mouse hippocampal neurons was observed by fluorescence microscopy (Figure 1). Although the axons were crowded with cargos, we were able to track a few individually. The center position (*X*) of each cargo was obtained from fluorescence images (Methods). Figure 2A shows a typical example time course of cargo position (Supporting Figures S1 and S2). We observed that cargo velocities changed over the course of several seconds. In particular, the constant velocity segments (CVSs) of the time courses (Figure 2A, square) with velocities greater than 100 nm/s and which last more than 0.5 s, were used for the following analyses on driving force (Figure 1), assuming that CVS fluctuation (Figure 2B) in one direction was affected minimally by other motors (the assumption is discussed in the Discussion section).

**Fig. 1.**
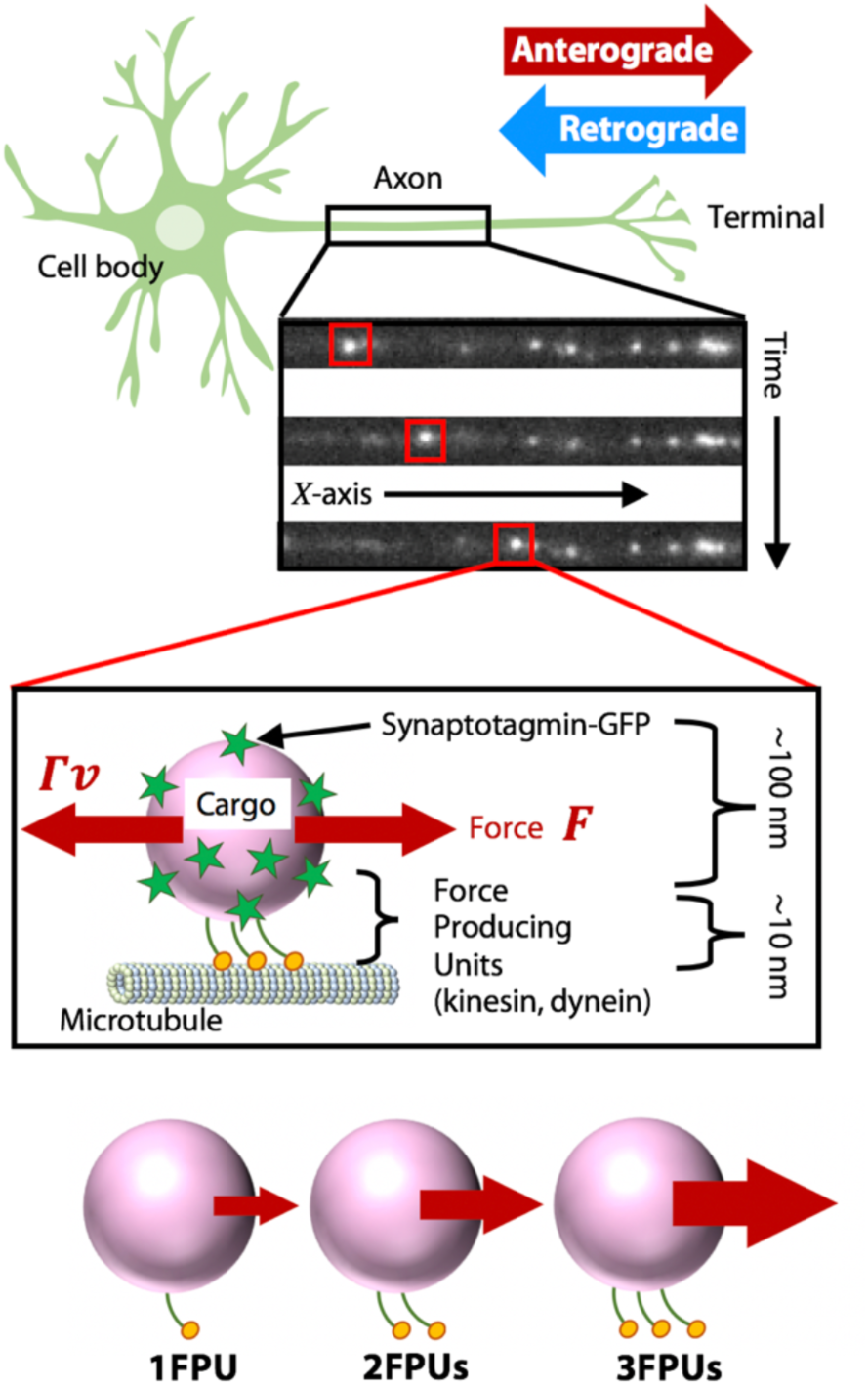
Synaptic cargo transport in the axon of a hippocampal neuron. Synaptic cargos (pink circles) labelled with GFP (green stars) undergo anterograde transport by kinesin and retrograde transport by dynein along microtubules (green cylinder). White dots in the fluorescence micrographs represent synaptic cargo. The force (*F*) is generated by the motor proteins and equivalent to the drag force, Γ_*v*_, at a constant velocity segment (CVS), where v and Γ represent the velocity and friction coefficient of moving cargo, respectively. Bottom: A single cargo transported by multiple force producing units (FPUs).

**Fig. 2.**
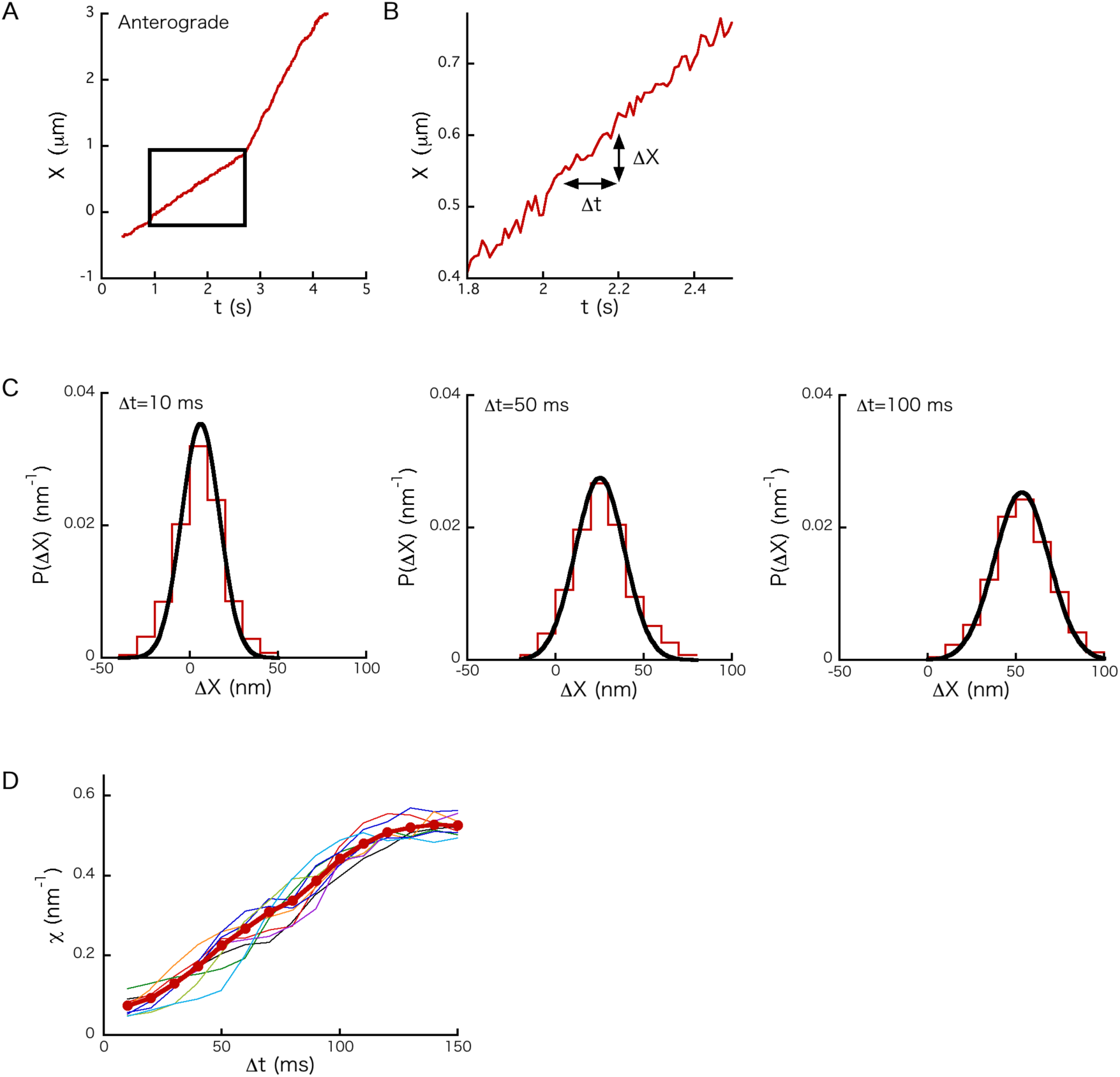
Fluctuation of position of a transported cargo. (A) Typical example of cargo position (*X*) over time. The circled time interval is a constant velocity segment (CVS). (B) Magnified view of the time course (Figure 2A) at a CVS. *Δ X* = *X* (*t* + *Δt*) − *X* (*t*). (C) Distribution *P*(*ΔX*) of *Δ X* for the cases in which *Δt* = 10 ms (left), 50 ms (middle), and 100 ms (right). *P*(*ΔX*) fit a Gaussian distribution (black curve). (D) Example of *χ* (*n*=1) for the CVS (Figure 2A) calculated using eqn (M3) plotted as a function of *Δt* (the thick curve). The thin curves represent *χ* calculated from different partial segments cut from the original CVS to estimate the error of *χ* in a boot-strapping manner (Methods). After relaxation time (about 100 ms), *χ* reached a constant value.

The fluctuation of a CVS is represented by the distribution *P*(Δ*X*), where Δ*X* = *X*(*t* + Δ*t*) − *X*(t) (Figure 2B). *P*(Δ*X*) is plotted when Δ*t* = 10 ms, 50 ms, and 100 ms (Figure 2C). The peak of *P*(Δ*X*) shifts to the right because of the directional motion of transported cargo, and variance increases due to the diffusional effect, as Δ*t* becomes larger. *P*(Δ*X*) was well described by a Gaussian function as well as those measured for other intracellular cargos (Hasegawa et al., 2019; Hayashi, Hasegawa, et al., 2018; Hayashi, Tsuchizawa, et al., 2018). Note that the abnormal diffusion property (i.e., *P*(Δ*X*) is not a Gaussian function) often reported for intercellular cargo transport(Fakhri et al., 2014; Posey, Blaisdell-Pijuan, Knoll, Saif, & Ahmed, 2018; Shin et al., 2019; Tabei et al., 2013) appeared upon analysis of the whole trajectory without dividing it into CVSs(Hayashi et al., 2013). One reason for the abnormality might be that the trajectory includes multiple velocity values.

### 2.3 Force index (*χ*) and quantification of FPUs transporting a cargo

Using *P*(Δ*X*) calculated in Figure 2C, an example force index *χ* (eqn (M3)) for the CVS was calculated as a function of Δ*t* as described in Methods (Figure 2D, thick curve). Over time, it converges to a constant value (Δ*t* > 100 ms). This relaxation time is characteristic for microscopic interactions, such as tethering interactions between the cargo and microtubules, collisions with other organelles and cytoskeletons, and ATP hydrolysis by motors(Hayashi, Tsuchizawa, et al., 2018). Then, we plotted *χ* for about 100 anterograde (Figure 3A, left) and retrograde (Figure 4A, left) moving cargos, respectively. Note that direction in motion was set to the positive direction when *χ* was calculated. The discreteness of *χ* was found with synaptic cargo transport and was observed previously for other cargo transports(Hasegawa et al., 2019; Hayashi, Hasegawa, et al., 2018; Hayashi, Tsuchizawa, et al., 2018). After applying a classification method (Methods), we found 6 clusters for both anterograde (Figure 3A) and retrograde (Figure 4A) transport. This indicates that we identified 6 FPUs when the discreteness of the force index *χ* represented the discreteness of transport force. Note that two trajectories of *χ* (∼2 nm^-1^) were excluded from cluster analysis for retrograde transport because the clustering analysis became difficult when such a small number of trajectories of *χ* were included (see Figure S4 for the trajectories). One candidate FPU is a kinesin dimer for anterograde transport, and four monomers of dynein have been implicated for retrograde transport by cryo-electron microscopy(Urnavicius et al., 2018). However, the constitution of one FPU is not elucidated, yet.

**Fig. 3.**
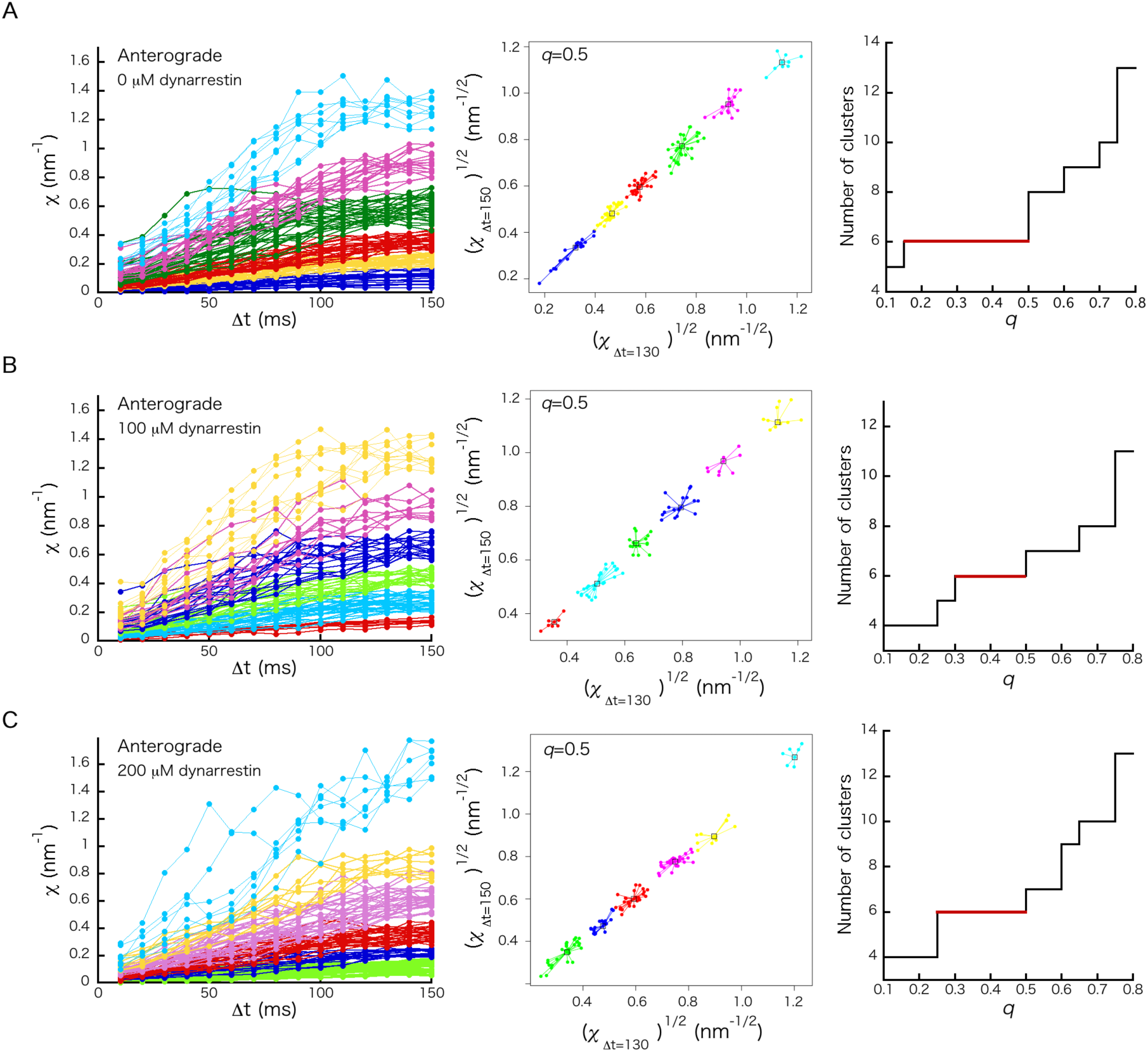
Force index χ for anterograde transport. *χ* as a function of *Δt* for *n* different cargos in cells treated with (A) 0 μM dynarrestin (*n* = 131), (B) 100 μM dynarrestin (*n* = 92), and (C) 200 μM dynarrestin (*n* = 119) (left panels). Each color denotes a cluster (i.e., FPU). The number of clusters was determined by clustering analysis (described in Methods) (middle panels). The number of clusters are displayed as a function of *q*, which is the sole parameter of the cluster analysis (right panels). The most probable cluster number was decided as the number valid for the wide range of *q*.

**Fig. 4.**
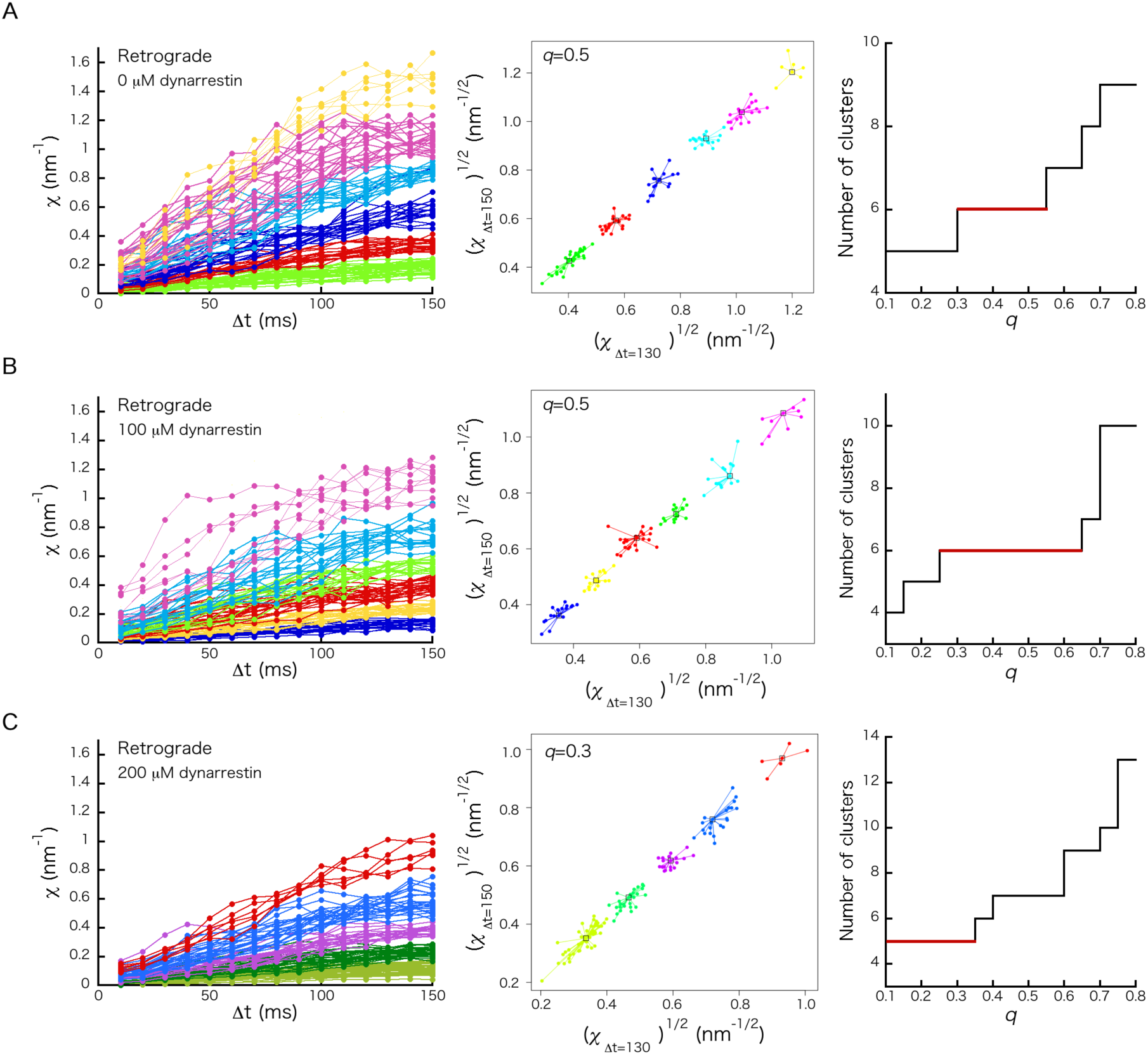
Force index *χ* for retrograde transport. *χ* as a function of *Δt* for *n* different cargos in cells treated with (A) 0 μM dynarrestin (*n* = 116), (B) 100 μM dynarrestin (*n* = 102), and (C) 200 μM dynarrestin (*n* = 123) (left panels). Each color denotes a cluster (i.e., FPU). The number of clusters was decided by clustering analysis (described in Methods) (middle panels). The right panels show the number of clusters as a function of *q*, which is the sole parameter of the cluster analysis. The most probable cluster number was decided as the number valid for the wide range of *q*.

### 2.4 Addition of the dynein inhibitor

We investigated the effect of the drug dynarrestin on synaptic cargo transport to study whether *χ* can detect the motor number change caused by the inhibitor. Dynarrestin inhibits ATPase activity of AAA+ ATPase superfamily, including dynein(Hoing et al., 2018). We treated neurons with 100 μM and 200 μM dynarrestin and plotted anterograde (Figure 3B, C) and retrograde (Figure 4B, C) transport. Changes in anterograde FPUs were small; retrograde FPUs showed larger decreases due to the dynarrestin treatment (Figure 5A).

**Fig. 5.**
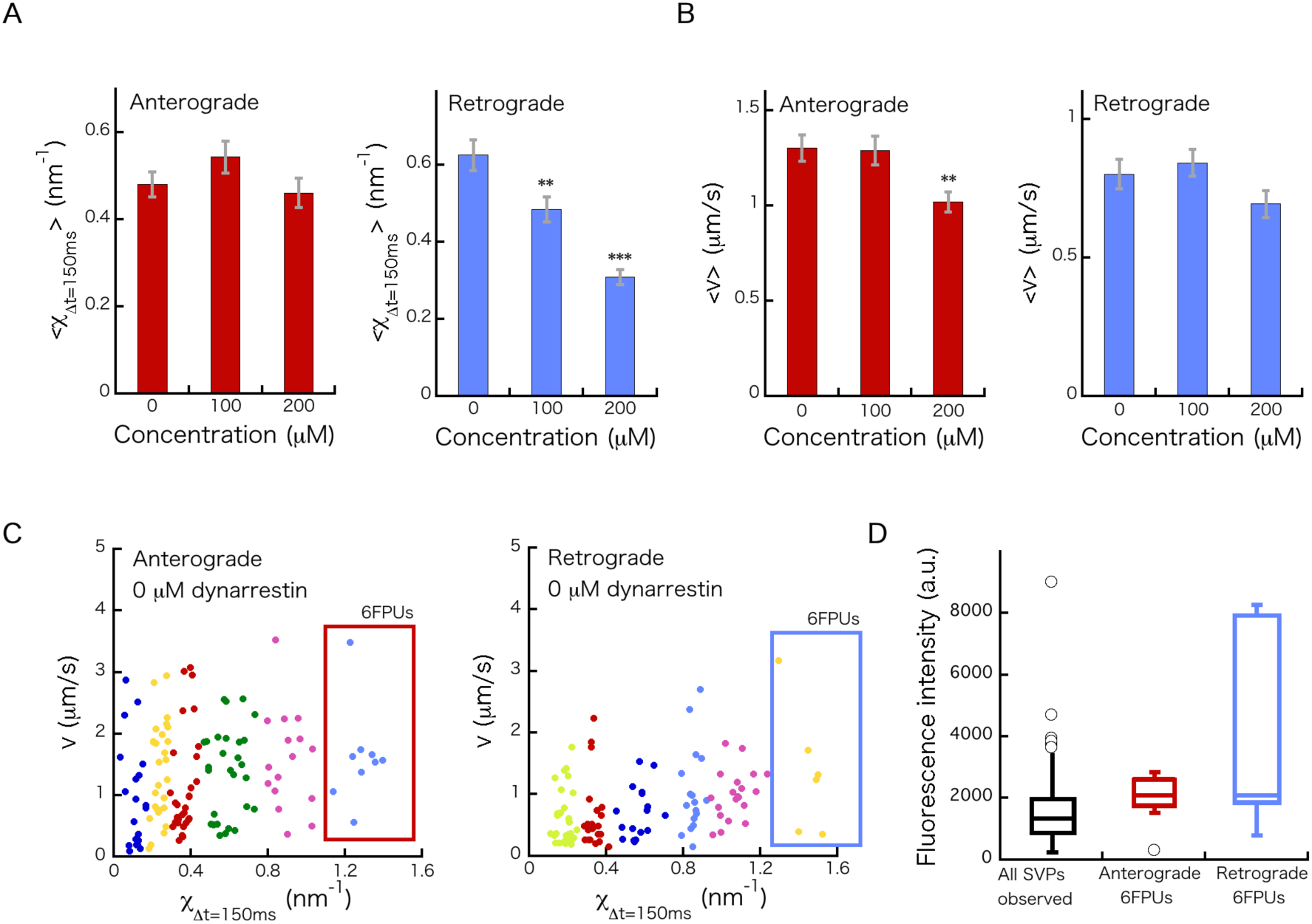
Relationship between *χ*, velocity, and fluorescence intensity. (A) Mean values of *χ* as a function of dynarrestin concentration for anterograde (left) and retrograde (right) transport. Error bars represent standard error (SE). (B) Mean velocity (*v*) at the CVSs as a function of dynarrestin concentration for anterograde (left) and retrograde (right) transport. Error bars represent standard error (SE). (C) Relationship between *χ* in the case *Δt* =150 ms and *v* in the absence of dynarrestin for the case of anterograde (left) and retrograde (right) transport. Each color of the dots denotes each cluster shown in Figure 3A and 4A, respectively. (C) Comparison among fluorescence intensities of all cargos observed, anterograde cargos belonging to six FPUs, and retrograde cargos belonging to six FPUs, respectively.

Compared with the large decrease in the number of retrograde FPUs (Figure 5A), the decrease in velocity (*v*) in the presence of dynarrestin was small (Figure 5B). Noting that *v* depends on cargo size (Supporting Figure S5) as well as the number of FPUs, this small change in velocity is explained by the relationship between *χ* and *v* (Figure 5B). Cargo with large FPUs did not always move with the greatest velocity because the fluorescence intensities of cargo, which represent the sizes of cargos, with 6 FPUs are greater than other cargos (Figure 5D). This indicates that the lack of large FPUs caused by the dynarrestin treatment contributed only slightly to the decrease in velocity.

In Figure 6, the mean values of *χ* (*Δt* = 150 ms) for each FPU were plotted as a function of dynarrestin concentration in terms of anterograde (left) and retrograde (right) transport. While anterograde transport changed very little upon dynarrestin treatment, there was a tendency for the retrograde mean values to decrease as the dynarrestin concentration increased. This trend may have been due to damage by the inhibitors to some of multiple dynein monomers constituting one retrograde FPU.

**Fig. 6.**
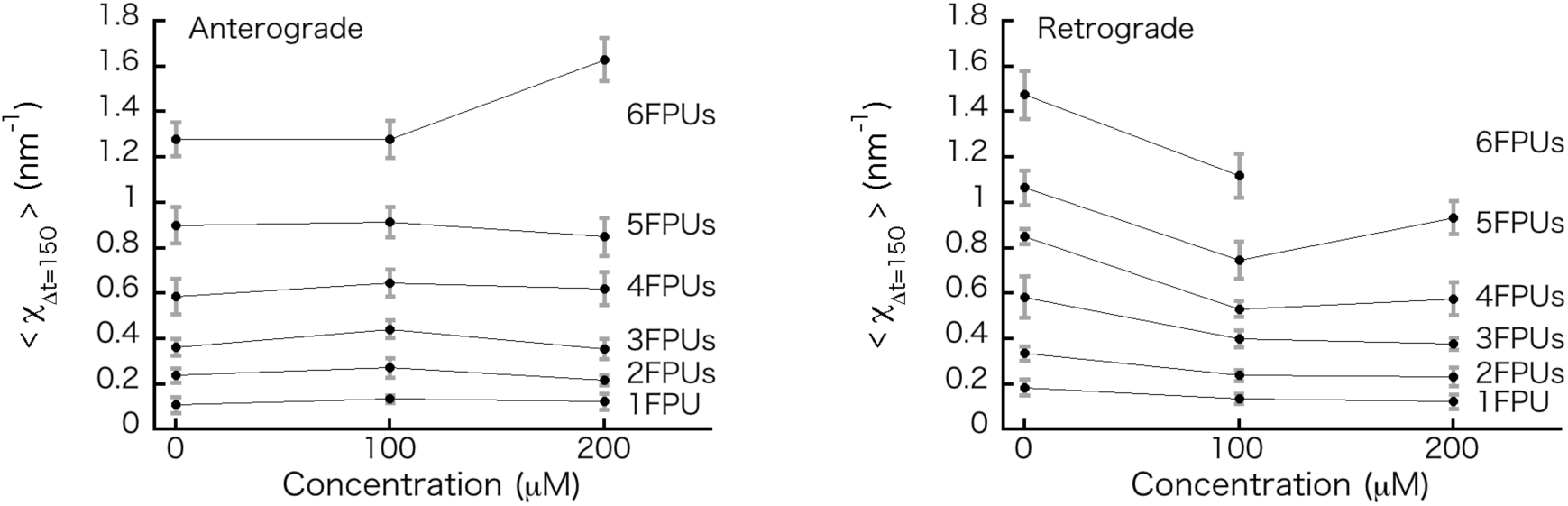
Mean values of *χ* (*Δt* = 150 ms) (⟨*χ*_*Δt* = 150_⟩) for each FPU. **⟨***χ* _***Δt* = 150**_**⟩** as a function of dynarrestin concentration in anterograde (left) and retrograde (right) transport. **⟨***χ* _***Δt* = 150**_**⟩** was calculated using the elements belonging to each FPU group. Error bars represent standard deviation (SD).

## 3. Discussion

In this study, we used fluorescence microscopy to observe synaptic cargo in the axons of mouse hippocampal neurons, in which cargo undergo anterograde transport by KIF1A and KIF1β and retrograde transport by cytoplasmic dynein (Figure 1). Given that cooperative activity by multiple motors contributes to stable and long-distance transport, and that FPU number represents an index of healthy axonal transport, we measured the force index *χ* (eqn (M3)) and identified 6 FPUs used for anterograde (Figure 3A) and retrograde (Figure 4A) transport. We then quantified the effect of the dynein inhibitor dynarrestin(Hoing et al., 2018) on cargo transport (Figure 5A). Given the difficulty of direct physical measurement in cells, we hope that application of this technique promotes further understanding of the mechanisms of intracellular cargo transport, including the roles of adaptor proteins connecting cargos and motors(Chiba et al., 2014). Below, two observations related to the main results are discussed.

Based on the fluctuation theorem(Ciliberto, Joubaud, & Petrosyan, 2010; Evans, Cohen, & Morriss, 1993), the relationship between *χ* and the driving force (*F*) is represented as

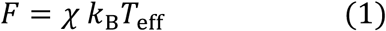

where *k*_B_ is the Boltzmann constant and *T*_eff_ is an effective temperature(Hasegawa et al., 2019; Hayashi, Tsuchizawa, et al., 2018), which is a generalized temperature in a non-equilibrium system(Crisanti & Ritort, 2003; Cugliandolo, 2011; Hayashi & Sasa, 2004). In our previous studies on intracellular cargo transport(Hasegawa et al., 2019; Hayashi, Hasegawa, et al., 2018; Hayashi, Tsuchizawa, et al., 2018), *T*_eff_ was not equal to the temperature of the environment (*T*), due to non-equilibrium effects. When one anterograde FPU corresponded to a kinesin dimer (which has a stall force of about 5 pN), *T*_eff_ was estimated to be about 10 times *k*_B_*T* for synaptic cargo transport. Note that when *T*_eff_ takes a different value for each moving cargo, the discreteness of the force index 3 disappears. Indeed, the constancy of *T*_eff_ was cross-checked with the relationship *χ ∝ v*, derived from eqn (1) and the Stokes’ relationship *F* = Γ*v* (Γ: friction coefficient), for the long time courses of the same cargos, in which Γ of the same cargo was considered constant during the time courses (Supporting Figure S1-S3). Finally, we consider the assumption that the CVS fluctuation (Figure 2B) in one direction was affected minimally by other motors. This assumption is not trivial, because the tug-of-war (TOW) motion between two motors(Gross, 2004; Muller, Klumpp, & Lipowsky, 2008; Soppina, Rai, Ramaiya, Barak, & Mallik, 2009; Welte, 2004), which could be an origin of this fluctuation, occurs during changes in direction. Because the mean value of the anterograde force index *χ* for each FPU changed minimally (Figure 6, left) along with decrease of the retrograde FPUs caused by the inhibitor (Figure 5A), this indicated that CVS fluctuation in one direction was not sensitive to the number of other motors. Thus, CVSs were considered separately from TOW segments(Soppina et al., 2009).

## 4. Methods

### 4.1 Primary culture of neurons and transfection

Primary culture of hippocampal neurons was performed using a previously described method with some modification (Bartlett & Banker, 1984; Chiba et al., 2014). The hippocampus of wild-type C57BL/6 mice (Japan SLC, Hamamatsu, Japan) at embryonic day 17 was dissected, and neurons were cultured in glass-bottom dishes (MatTek, Ashland, MA, USA) with culture medium (NbActiv4; BrainBits, Springfield, USA), as described previously(Chiba et al., 2014). After culture for 4 to 7 days, neurons were transfected with the plasmid vector for GFP-fused synaptotagmin(Niwa et al., 2013) using the calcium phosphate method (Takara Bio, CA, USA). All animal experiments were conducted in compliance with a protocol approved by the Institutional Animal Care and Use Committee, Tohoku University.

### 4.2 Dynarrestin treatment and fluorescence microscopy

Dynarrestin (Tocris Bioscience, Bristol, UK) was added to a culture dish with neurons one day post-transfection (final concentration of dynarrestin was 100 or 200 μM), and the dish was incubated for 30 min prior to fluorescence observation, then washed with Hank’s balanced salt solution (Thermo Fisher Scientific, Waltham, MA, USA) containing 100 μM or 200 μM dynarrestin.

Cargo movement was observed with a fluorescence microscope (IX83; Olympus, Tokyo, Japan) equipped with a heating plate (CU-201; Live Cell Instrument, Seoul, Korea) maintained at 37 °C for 45 min. For this observation, B27-supplement (Thermo Fisher Scientific) and 150 μM 2-mercaptoethanol (Wako Chemical Co., Tokyo, Japan) were added to the medium. Images were obtained with a 150× objective lens (UApoN 150×/1.45; Olympus) and an sCMOS camera (OLCA-Flash4.0 v.2.0; Hamamastu Photonics, Hamamatsu, Japan) at 100 frames/s. The center position of each cargo vesicle was determined from the recorded images using custom software written in LabVIEW 2013 (National Instruments, Austin, TX, USA), as described previously(Hayashi, Hasegawa, et al., 2018).

The primary culture of neurons was repeated seven times, and data were collected from six to eight preparations for each dynarrestin concentration. To examine anterograde and retrograde transport, we investigated 131 and 116 cargos from 50 and 39 different cells, respectively, for untreated cells, 92 and 102 cargos from 35 and 37 different cells, respectively, treated with 100 μM dynarrestin, and 119 and 123 cargos from 43 and 41 different cells, respectively, treated with 200 μM dynarrestin, respectively. The cells used for observation were chosen randomly after visual inspection.

The accuracy of positional measurements was verified with beads (300-nm latex beads; Polyscience, Niles, IL, USA) of similar size compared to the synaptic cargos. The standard deviation of bead position tightly attached to the glass surface was 8.3 ± 1.2 nm (*n* = 4 different beads).

### 4.3 Calculation of *χ*

The force index *χ* introduced in our previous studies(Hasegawa et al., 2019; Hayashi, Hasegawa, et al., 2018; Hayashi, Tsuchizawa, et al., 2018) was originally defined using the idea of the fluctuation theorem as follows:

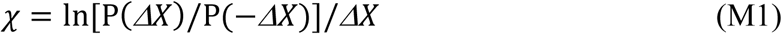

from the distribution *P*(Δ*X*) of the displacement *ΔX* = *X*(*t* + *Δt*) − *X*(*t*) (Figure 2B). *P*(Δ*X*) was fitted with a Gaussian function (Figure 2C):

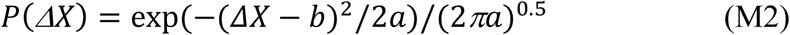

where the fitting parameters *a* and *b* correspond to the variance and mean of the distribution, respectively. By inserting eqn (M2) into eqn (M1), we obtain:

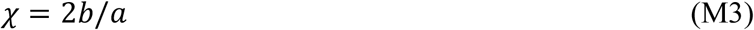

Equation (M3) is equivalent to the equation *χ* = *v/D* when the diffusion coefficient *D* is defined as *D* = *a/*2*Δt*. The error of *χ* was estimated at 10% based on the boot-strapping method(Hasegawa et al., 2019; Hayashi, Hasegawa, et al., 2018) that *χ* calculated from different partial segments cut from the original CVS (n = 10), whose length was 50% of the original segment (Figure 2D, thin curves).

A smoothing filter was applied to the values of *χ* to reduce variation in the raw data for *χ* as a function of Δ*t*. We used the following simple averaging filter:

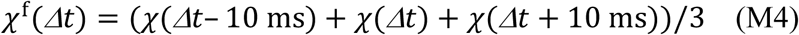

Note that *χ* ^f^ (*Δt*) = (*χ* (*Δt*) + *χ* (*Δt* + 10 ms))/2 for *Δt* = 10 ms and *χ*^f^(*Δt*) = (*χ* (*Δt* − 10 ms) + *χ* (*Δt*))/2 for *Δt* = 150 ms. The software used to calculate *χ* is available from a website(“It’ll be provided via GitHub.,”).

### 4.4 Classification of *χ*-*Δt* plots

Affinity propagation(Bodenhofer, Kothmeier, & Hochreiter, 2011; Frey & Dueck, 2007), an exemplar-based clustering method that does not require the number of clusters, was adoptedtoclusterthetwo-dimensionaldata 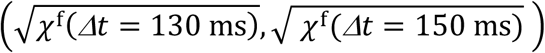. The square root of *χ* was used to apply the cluster method to make the distance between adjoining clusters constant because the value of 3 was observed to exponentially increase as the number of FPUs increased, although the reason for this was not understood. The two-dimensional data(*χ* ^f^ (*Δt* = 130 ms), *χ* ^f^ (*Δt* = 150 ms)) was also available for the clustering, though was less stable (the decrease of the values of retrograde *χ* in the presence of dynarrestin (Figure 6, right) made the clustering analysis difficult).

The method was applied using the ‘APCluster’ package in R(“R Core Team. R: Alanguage and environment for statistical computinh. R Foundation for Statistical Computing, Vienna, Austria. URL https://www.R-project.org/,” 2018). The clustering was stable for the wide range of values for the sole parameter (*q*) of the affinity propagation analysis.

### 4.5 Statistical test

Student’s t-test was applied by using the function (t.test()) of R software(“R Core Team. R: Alanguage and environment for statistical computinh. R Foundation for Statistical Computing, Vienna, Austria. URL https://www.R-project.org/,” 2018) to the data in Figure 5A and 5B. **p < 0.01 and ***p < 0.001.

## Acknowledgements

We acknowledge Dr. K. Chiba for providing the method used to culture hippocampal neurons, S. Hasegawa for assistance with the particle-tracking software, and K. Nagino for experiment support. We would like to thank Edita<u>g</u>e (www.editage.com) for English language editing. This work was supported by JST PRESTO (grant No. JPMJPR1877), AMED PRIME (grant No. JP18gm5810009), and JSPS KAKENHI (grant No. 17H03659) to K. H., as well as JSPS KAKENHI (grant Nos. 17H05010 and 19H04738) to S. N.

## Conflict of interests

The authors declare no conflict of interests.

## Author Contributions

K. H. and M. G. M performed the experiments. K. H. analyzed the data and wrote the paper with contributions from S. N.

## Supporting Information

Supporting Figures S1-S5

**Fig. S1.**
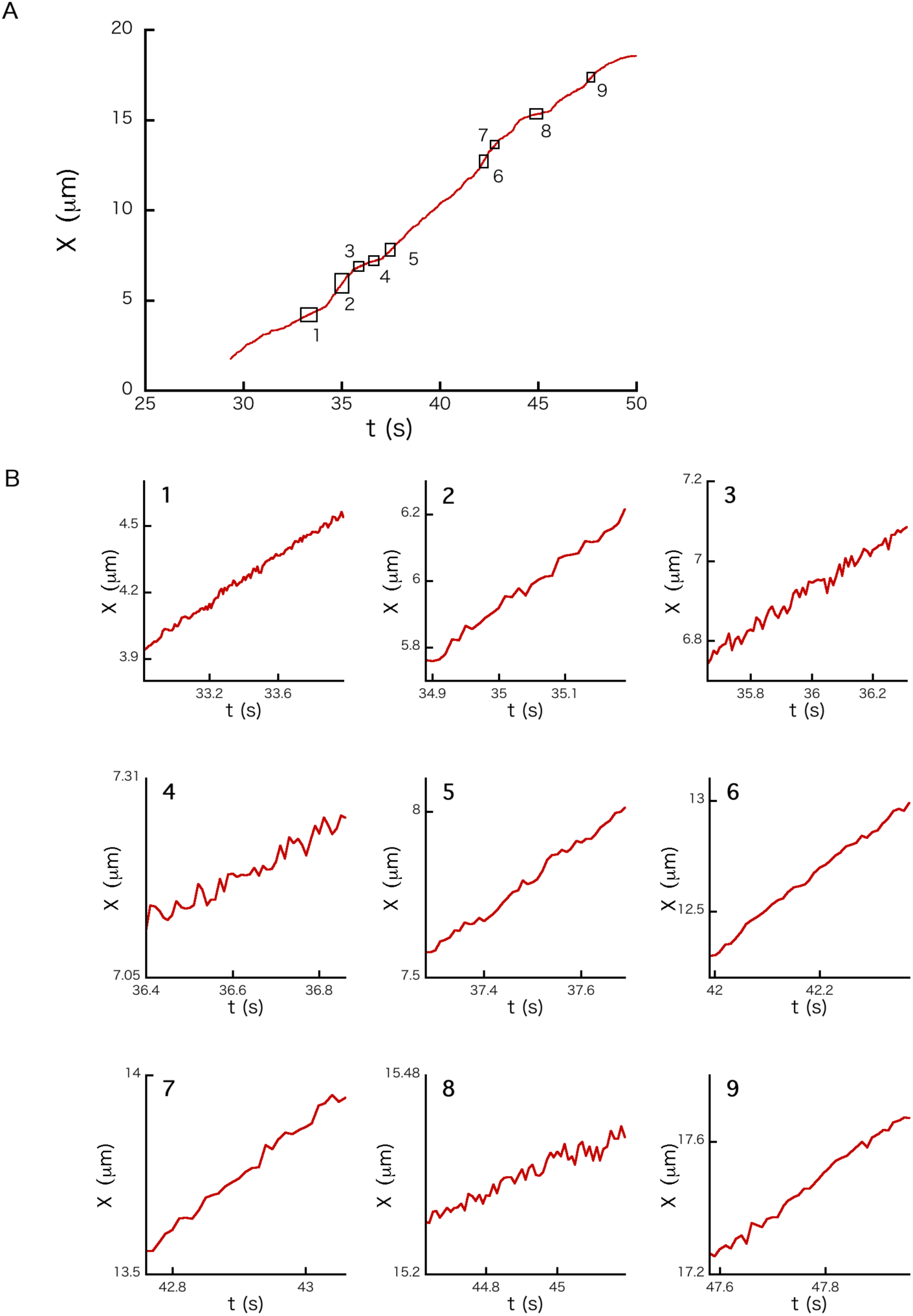
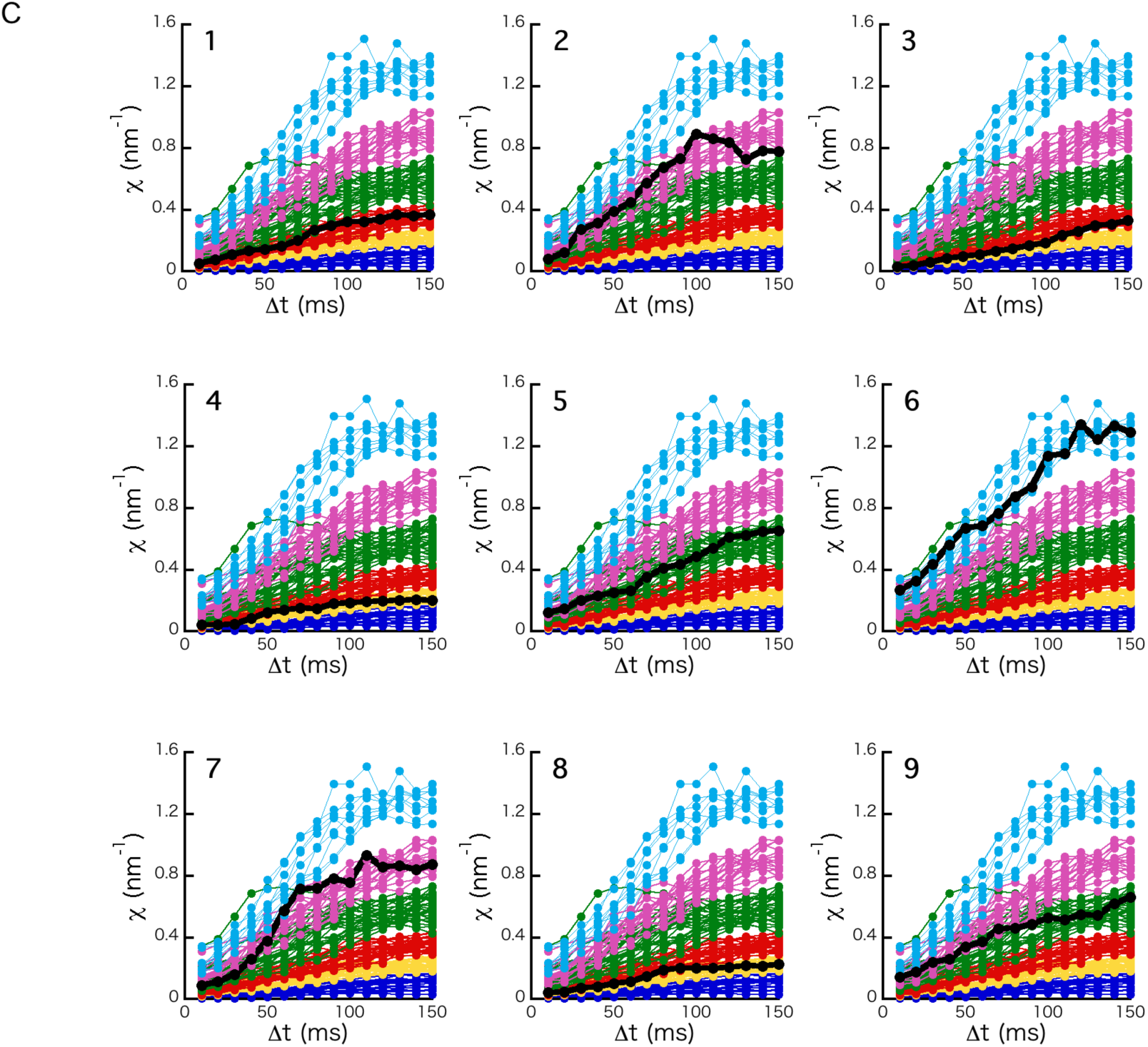
Analysis of a long time course of the position (*X*) of a cargo anterogradely moving. (A) Example long time course of *X* in the absence of dynarrestin. The circled time intervals of *X* are constant velocity segments (CVSs). (B) Magnified views of the CVSs shown in Figure S1A. (C) *χ* as a function of *Δt* for each CVS (thick black curves). Note that direction in motion was set to the positive direction when *χ* was calculated. The background thin curves are originally shown in Figure 3A of the main text.

**Fig. S2.**
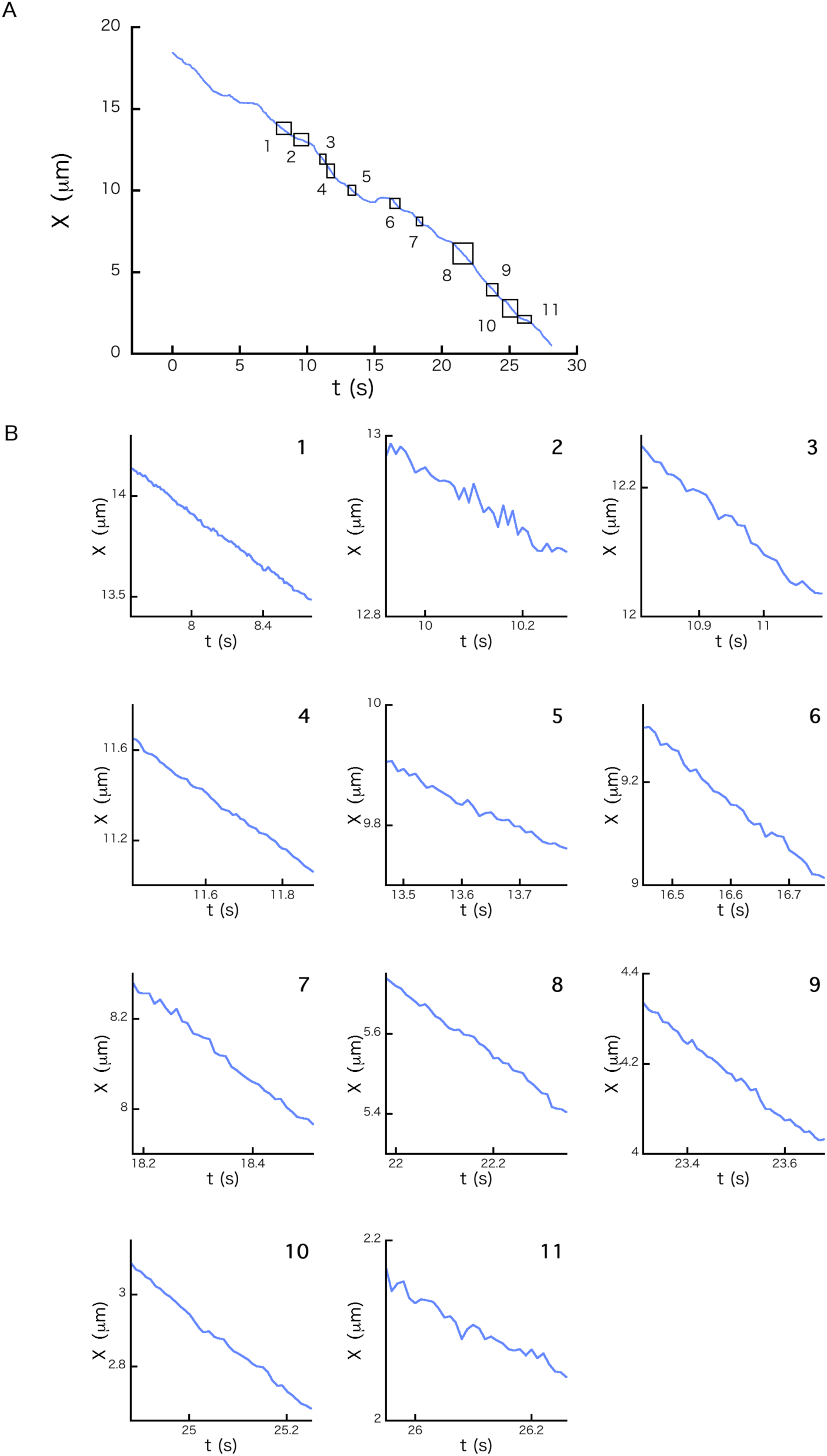
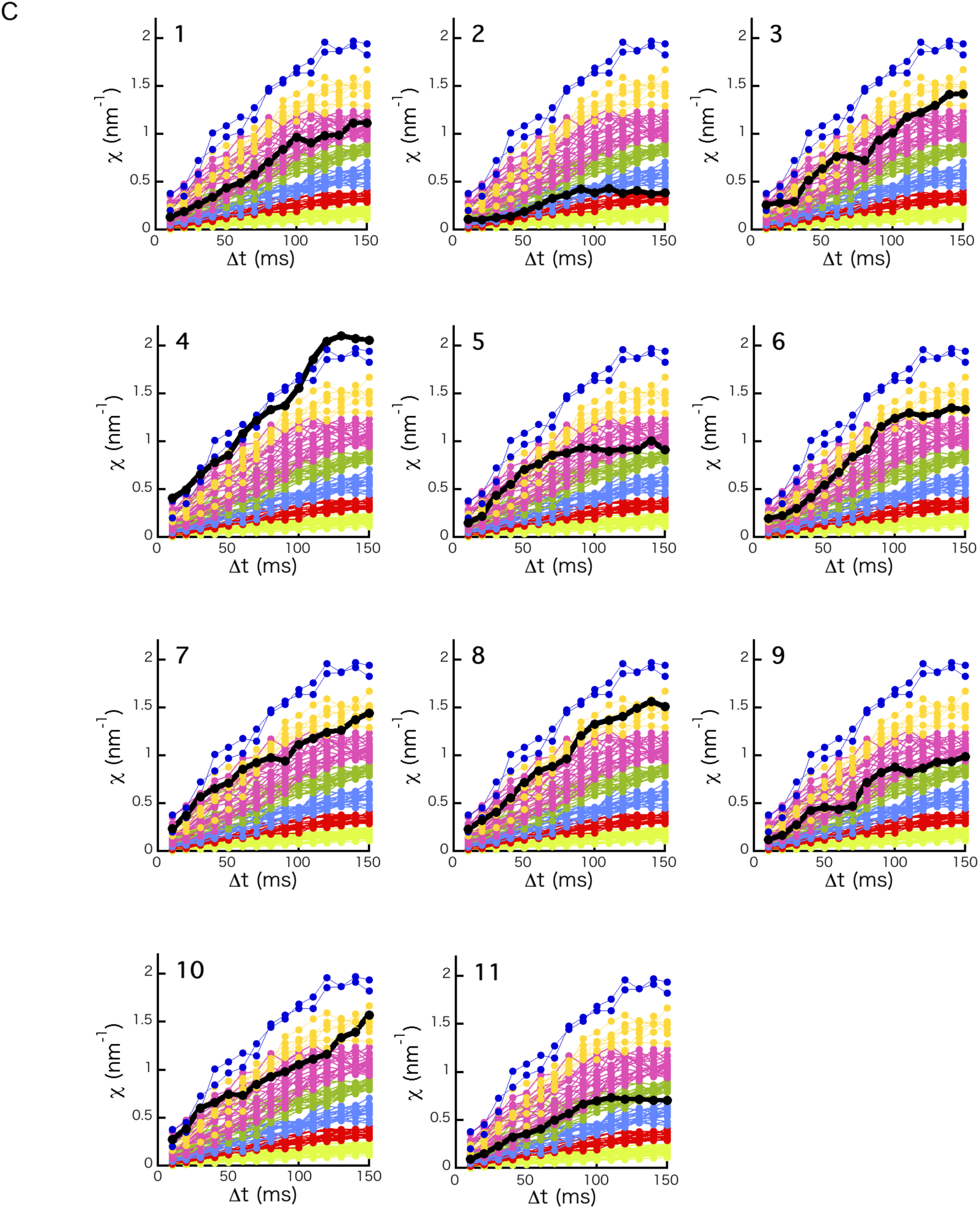
Analysis of a long time course of the position (*X*) of a cargo retrogradely moving. (A) Example long time course of *X* in the absence of dynarrestin. The circled time intervals of *X* are constant velocity segments (CVSs). (B) Magnified views of the CVSs shown in Figure S2A. (C) *χ* as a function of *Δt* for each CVS (thick black curves). Note that direction in motion was set to the positive *X* direction when *χ* was calculated. The background thin curves are originally shown in Figure 4A of the main text.

**Fig. S3.**
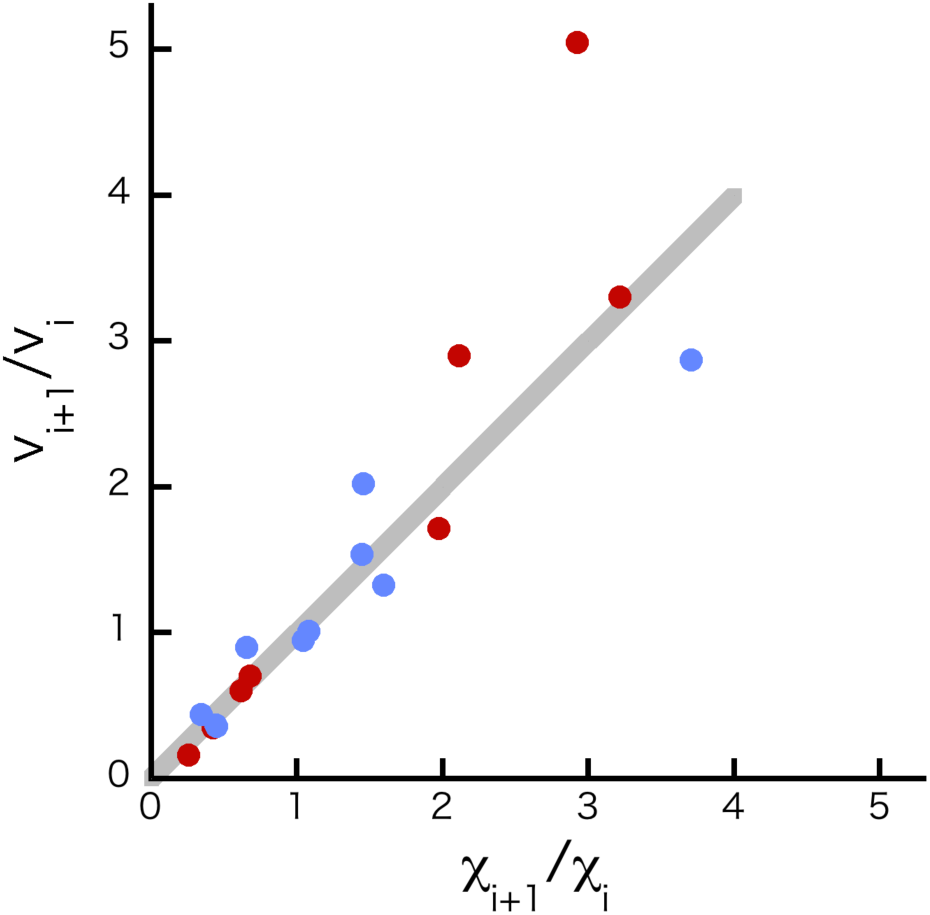
Relationship between *χ* and *v* for the same cargo. *χ*_i+1_/*χ*_i_ as a function of *v*_i+1_/*v*_i_ in the cases of the anterograde CVSs shown in Figure S1B (red), and the retrograde CVSs shown in Figure S2B (blue). *i* represents a segment number of the graph in Figure S1B and S2B. The grey line represents *χ*_i+1_/*χ*_I_ = *v*_i+1_/*v*_i_.

**Fig. S4.**
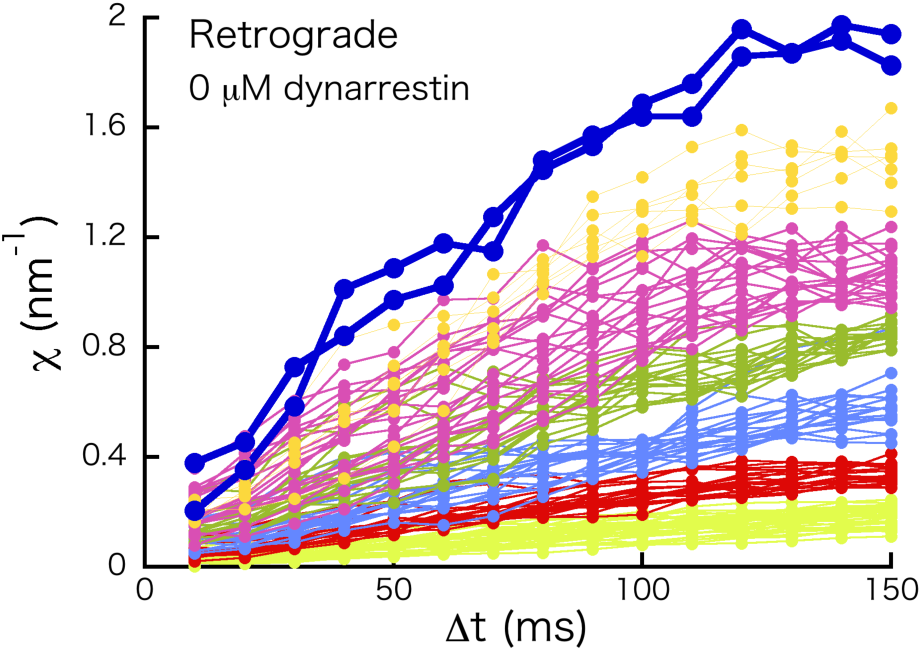
Force index *χ* for retrograde transport including *χ*∼2 nm^-1^ (thick navy curves). The thin curves are originally shown in Figure 4A (left).

**Fig. S5.**
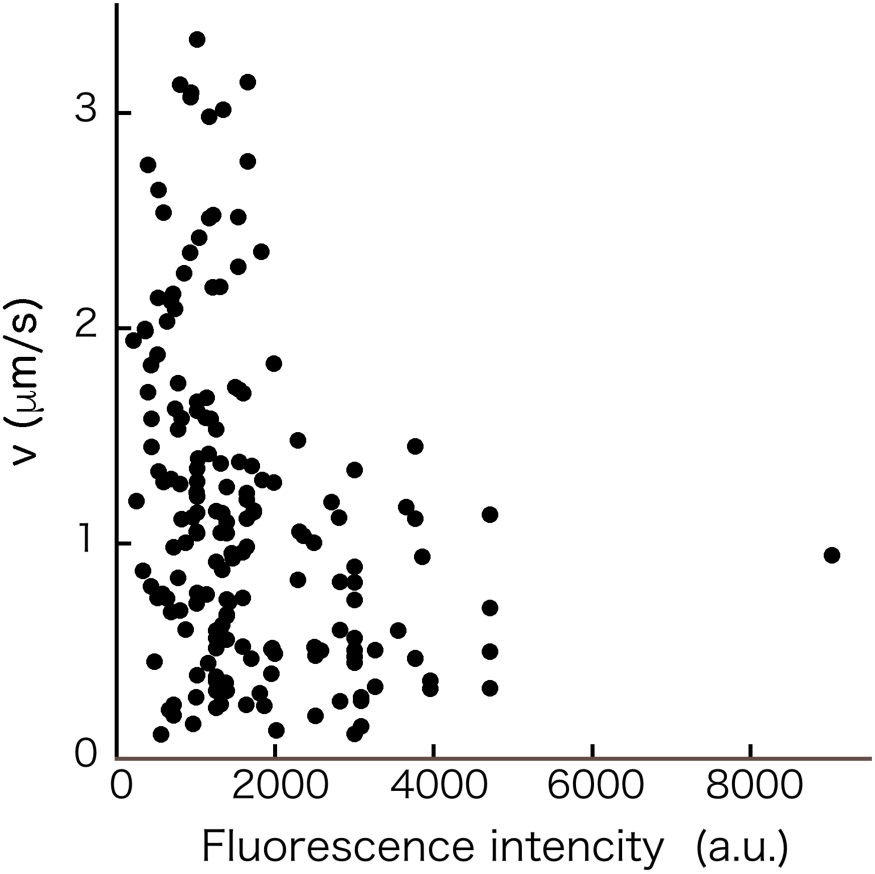
Velocity *v* of a constant velocity Segment (CVS) as a function of the fluorescence intensity of a synaptic cargo in the absence of dynarrestin. Anterograde and retrograde data were not separated.

## Notes

### Competing Interest Statement

The authors have declared no competing interest.

### Summary of Updates

The results obtained by using ciliobrevin were replaced with those obtained by using dynarrestin.

